# Physiopathological changes of ferritin mRNA density and distribution in hippocampal astrocytes in the mouse brain

**DOI:** 10.1101/2022.04.01.486678

**Authors:** Romain Tortuyaux, Marc Oudart, Noémie Mazaré, Philippe Mailly, Jean-Christophe Deschemin, Sophie Vaulont, Carole Escartin, Martine Cohen-Salmon

**Affiliations:** Physiology and Physiopathology of the Gliovascular Unit Research Group, Center for Interdisciplinary Research in Biology (CIRB), College de France, CNRS Unité Mixte de Recherche 724, Inserm Unité 1050, Labex Memolife, Université PSL, 75005 Paris, France; École doctorale Cerveau Cognition Comportement “ED3C” N°158, Pierre and Marie Curie University, 75005 Paris, France; CHU Lille, Intensive Care Unit, Lille, France; Orion Imaging Facility, Center for Interdisciplinary Research in Biology (CIRB), College de France, CNRS Unité Mixte de Recherche 724, INSERM Unité 1050, Labex Memolife, PSL Research University, Paris, France; Université de Paris Cité, CNRS, INSERM, Institut Cochin, F-75014 Paris, France; Université Paris-Saclay, CEA, CNRS, MIRCen, Laboratoire des Maladies Neurodégénératives, Fontenay-aux-Roses, France

## Abstract

Astrocytes are thought to play a crucial role in brain iron homeostasis. How they accomplish this regulation *in vivo* remains unclear. In a recent transcriptomic analysis, we showed that polysomal Ftl1 and Fth1 mRNAs, encoding the ferritin light (Ftl) and heavy (Fth) chains that assemble into ferritin, a critical complex for iron storage and reduction, are enriched in perisynaptic astrocytic processes as compared to astrocytic soma. These data suggested that ferritin translation plays a specific role at the perisynaptic astrocytic interface and is tighly regulated by local translation. Here, we used our recently described *AstroDot* 3D *in situ* methodology to study the density and localization of ferritin mRNAs in astrocytes in the hippocampus in three different contexts in which local or systemic iron overload has been documented: ageing, the hepcidin knock-out mouse model of hemochromatosis and the APP/PS1dE9 mouse model of Alzheimer’s disease (AD). Our results showed that in wild type mice, Fth1 mRNA density was higher than Ftl1 and that both mRNAs were mostly distributed in astrocyte fine processes. Ageing and absence of hepcidin caused an increased Fth1/Ftl1 ratio in astrocytes and in the case of ageing, led to a redistribution of Fth1 mRNAs in astrocytic fine processes. In contrast, in AD mice, we observed a lower Fth1/Ftl1 ratio and Fth1 mRNAs became more somatic. Hence, we propose that regulation of ferritin mRNA density and distribution in astrocytes regulates iron homeostasis in physiology and pathophysiology.

## Introduction

Iron is a component of heme groups and iron-sulfur clusters in proteins and is therefore essential for oxygen consumption, ATP production and a large variety of enzymatic reactions (Connor & Menzies, 1995). Iron homeostasis is tightly regulated since unbound iron has a toxic redox activity and iron excess or deficiency compromise cell viability. Several proteins work together to maintain iron homeostasis (Hentze, Muckenthaler, Galy, & Camaschella, 2010). Transferrin (Tf) is the most important iron carrier molecule. It is mainly synthesized in the liver, secreted in the blood and has the capacity to bind 2 atoms of ferric ion. Iron-loaded Tf interacts with the Tf receptor (TfR) on the cell surface and is internalized by endocytosis. In endosomes, ferric iron is reduced to ferrous iron by ferrous iron metallo-reductases and transported to the cytosol by the divalent metal transporter 1 (DMT1), while Tf is recycled back to the membrane. Free iron, although potentially highly toxic, can also be directly secreted by cells through ferroportin, also known as metal transport protein 1 (MTP1) (Atanasiu, Manolescu, & Stoian, 2007; Nemeth & Ganz, 2006). The activity of ferroportin is modulated by hepcidin, a small circulating peptide produced primarily in the liver. Hepcidin acts by binding to ferroportin, leading to its occlusion or degradation, thereby limiting cellular iron efflux. Ferritin is the primary intracellular iron-storage protein and is also essential for keeping iron in a soluble non-toxic form (Finazzi & Arosio, 2014). It consists in a 24 subunit heteropolymer composed by heavy (Fth encoded by Fth1) and light (Ftl encoded by Ftl1) chains. Fth has a ferroxidase activity converting soluble ferrous into ferric redox-inactive iron. Ferritin is able to bind over 4,500 atoms of iron (Harrison & Arosio, 1996). Due to its ability to sequester and reduce iron, ferritin is an important antioxidant protein.

Iron is essential to the brain physiology. Iron deficiency leads to learning and memory impairment (Lozoff et al., 2006), while its excess is associated with ageing and neurodegenerative disorders including Alzheimer’s disease (AD), Parkinson’s disease and multiple sclerosis (Belaidi & Bush, 2016; Liu, Liang, & Soong, 2019; Molina-Holgado, Hider, Gaeta, Williams, & Francis, 2007; Stephenson, Nathoo, Mahjoub, Dunn, & Yong, 2014; Zecca, Youdim, Riederer, Connor, & Crichton, 2004). Iron is also essential to oligodendrocyte differentiation and functions (Cheli, Correale, Paez, & Pasquini, 2020; Cheli et al., 2021; Stephenson et al., 2014). Thus, understanding how brain cells regulate iron levels is critical and the molecular mechanisms behind these regulations are potential therapeutic targets to control iron imbalance in the brain.

Astrocytes, an important glial cell type, are thought to be crucial actors in this matter (Cheli et al., 2020; Cheli et al., 2021; Dringen, Bishop, Koeppe, Dang, & Robinson, 2007; Hohnholt & Dringen, 2013). Astrocytes are highly polarized cells interfacing neurons and the blood vessels. Astrocytic perivascular processes (PvAPs) or endfeet entirely cover brain vessels and participate in the regulation of several vascular functions, in particular blood brain barrier (BBB) integrity (Abbott, Ronnback, & Hansson, 2006; Alvarez, Katayama, & Prat, 2013; Cohen-Salmon et al., 2021). Astrocytic perisynaptic processes (PAP) on the other hand regulate neurotransmission (Dallerac, Zapata, & Rouach, 2018; Perea, Navarrete, & Araque, 2009). *In vitro*, astrocytes have been shown to express most proteins involved in iron homeostasis. Hepcidin was detected in cortical and hippocampal astrocytes by immunofluorescence (Raha-Chowdhury et al., 2015; Zhang et al., 2020). DMT1 was found in PvAPs indicating that astrocytes can potentially uptake iron across the BBB (Wang, Ong, & Connor, 2002). Importantly, several studies pointed out the potential influence of astrocytes in handling iron homeostasis in aging and neuropathological contexts (Cheli et al., 2020; Dringen et al., 2007; Hohnholt & Dringen, 2013).

In a recent transcriptomic analysis, we demonstrated that both Fth1 and Ftl1 polysomal mRNAs are strongly enriched in hippocampal PAPs, suggesting that ferritin-mediated iron storage is prevalent in distal astrocytic processes such as PAPs (Mazare et al., 2020). Here, we aimed to further characterize Ftl1 and Fth1 mRNA distribution in hippocampal astrocytes of wild type (WT) mice as well as in conditions of impaired iron homeostasis: ageing, hepcidin-deficiency model of systemic iron overload (hemochromatosis) and AD.

## Materials and Methods

### Animals

All animal experiments were carried out in compliance with the European Directive 2010/63/EU on the protection of animals used for scientific purposes and the guidelines issued by the French National Animal Care and Use Committee. All mice were maintained on a C57BL6 genetic background. We carried out our study on 6-month-old hepcidin deleted mice (*Hepc1*-/-) (Lesbordes-Brion et al., 2006), 10-month-old APP/PS1dE9 mice (https://www.jax.org/strain/005864) (Jankowsky et al., 2004) and 6 and 10-month-old C57/Bl6 (WT) mice. Only females were included in this study except for iron dosage which was done on males.

### Brain slice preparation

Mice were anesthetized with a mix of ketamine/xylazine (140 and 8 mg/kg, respectively, i.p.) and killed by transcardiac perfusion with 1X phosphate-buffered saline (PBS)/4% paraformaldehyde (PFA). The brains were removed and immersed in 4% PFA overnight at 4°C. The PFA solution was replaced with 15% sucrose for 24 h at 4°C and, lastly, by 30% sucrose for 24 h at 4°C. The brains were cut into 40 µm-thick coronal sections using a Leitz microtome (1400). Sections were stored at −20°C in a cryoprotectant solution (30% glycerol and 30% ethylene glycol in 1X PBS).

### Fluorescent *in situ* hybridization and immunostaining

All procedures are detailed in (Oudart et al., 2020). FISH was performed using the RNAscope^®^ Multiplex Fluorescent Reagent Kit v2 (Advanced Cell Diagnostics Inc.) and specific probes (**Table S1**), according to the manufacturer’s instructions. Following the FISH procedure, slides were incubated with a blocking solution (0.2% normal goat serum, 0.375% Triton X-100, and 1 mg.mL^-1^ bovine serum albumin in 1X PBS) for 1 hour at room temperature (RT), incubated with an antibody against Glial fibrillary acidic protein (GFAP) overnight at 4°C (**Table S1**), rinsed three times with 1X PBS, and incubated with the secondary antibody (**Table S1**) for 2h at RT. Lastly, the slides were washed three times in 1X PBS and mounted in Fluoromount-G^®^ and DAPI (Southern Biotech).

### Imaging

Images were acquired using a Yokogawa W1 Spinning Disk confocal microscope (Zeiss) with a 63x oil objective (1.4 numerical aperture). The imaging conditions and acquisition parameters were the same for all slides. The experimental point spread function (PSF) was obtained using carboxylate microsphere beads (diameter: 170 nm; Invitrogen/ThermoFisher Corp.). Except for DAPI, all channels were deconvoluted with Huygens Essential software (version 19.04, Scientific Volume Imaging, The Netherlands; http://svi.nl), using the classic maximum likelihood estimation algorithm and a signal-to-noise ratio of 50 (for the immunofluorescence channel) or 20 (for the FISH channel), a quality change threshold of 0.01, and 150 iterations at most.

### FISH quantification by *AstroDot* and *AstroStat*

All procedures are detailed in (Oudart et al., 2020). Briefly, *AstroDot* is an ImageJ plug-in analyzing the localization of mRNAs at the level of astrocyte intermediate filaments immunolabelled against GFAP in soma, large and fine processes. Previous to the analysis, regions of interest (ROI) corresponding to individual astrocytes are selected manually by assessing the GFAP and DAPI (nuclei) staining on the Z projection of image stacks and by defining the stack of confocal planes.

#### FISH dot classification

A dot in the immunofluorescence background: mean GFAP immunofluorescence intensity ≤ bgThreshold; distance to the boundary of the nucleus > 2 µm. This type of dot was coloured in red and considered outside astrocytes.

A dot in a fine process: mean GFAP immunofluorescence intensity >bgThreshold; distance to the boundary of the nucleus >2 µm; astrocyte process diameter < step in the z calibration (0.3 µm). This type of dot was coloured in yellow.

A dot in a large process: mean GFAP immunofluorescence intensity >bgThreshold; distance to the boundary of the nucleus >2 µm; astrocyte process diameter > step in the z calibration (0.3 µm); or a dot in the soma if the distance to the boundary of the nucleus ≤2 µm. This type of dot was coloured in green.

### Brain iron dosage

Brain iron content was determined by acid digestion of 6-month-old WT and *Hepc1 -/-* dissected brains as described by Torrance and Bothwell (Torrance & Bothwell, 1980), followed by iron content determinations with a colorimetric assay on the Olympus AU400 automat.

### Astrodot statistics

Each analysis was carried out on groups of 3 mice. We used *AstroStat* an R package to analyse *AstroDot* data (Oudart et al., 2020) (R Core Team (2021). R: A language and environment for statistical computing. R Foundation for Statistical Computing, Vienna, Austria (https://www.r-project.org/). We used the Student’s t-test for normally distributed data and equal variances; the Welch-Satterthwaite’s test for a normal data distribution and unequal variances; the Wilcoxon’s test for non-normally distributed data. Histograms were generated using the ggplot2 package (Wickham H (2016). ggplot2: Elegant Graphics for Data Analysis. Springer-Verlag New York. ISBN 978-3-319-24277-4). The threshold for statistical significance was p < 0.05.

## Results

### In WT mice, ferritin mRNAs are preferentially distributed in fine hippocampal astrocytic processes and Fth1 is more abundant than Ftl1

We performed FISH experiments on 6-month-old WT whole hippocampal sections in order to detect Fth1 and Ftl1 mRNAs coding for the heavy and light ferritin chain in the CA1 region (**Fig. 1**). Samples were co-immunostained for the astrocyte-specific intermediate filament glial fibrillary acidic protein (GFAP) (**Fig. 1A**). FISH dots appeared to be distributed throughout astrocyte soma and processes (**Fig. 1A**). To characterize mRNA distribution, we used our recently developed *AstroDot* ImageJ plugin, which analyzes in 3D the localization of mRNAs at the level of GFAP-immunolabelled astrocyte somata, large and fine processes (Oudart et al., 2020) (**Fig. 1B, C)**. Individual astrocytes were selected manually by delineating the 3D astrocytic GFAP+ domain and their process diameter was calculated (see Materials and Methods) (**Fig. 1B**). Processes with a diameter greater than the minimum distance between two confocal planes (0.3 µm, in the present case), were defined as “large”, and those with a smaller diameter were defined as “fine”. The localization of mRNA FISH dots on GFAP+ processes was then analyzed. They appeared in green if they localized on astrocyte large GFAP+ processes and soma, or in yellow if they localized on astrocyte fine GFAP+ processes (**Fig. 1C**). Since ferritin is not exclusively expressed by astrocytes, we excluded from our analysis FISH dots not localized on GFAP+ processes, as they could originate from other brain cells (**Fig. S1**). The density and distribution of ferritin mRNAs in astrocytes was determined (**Fig. 1D, E**). We observed that Fth1 was expressed at higher levels than Ftl1 (8-fold) (**Fig. 1D, E**). Both mRNAs were mainly present in astrocytic processes and among them, in fine processes **(Fig. 1E)**. Overall, these results indicate that ferritin mRNAs are preferentially distributed in distal astrocyte compartments and that Fth1 is more abundant than Ftl1 mRNA.

**Figure 1.**
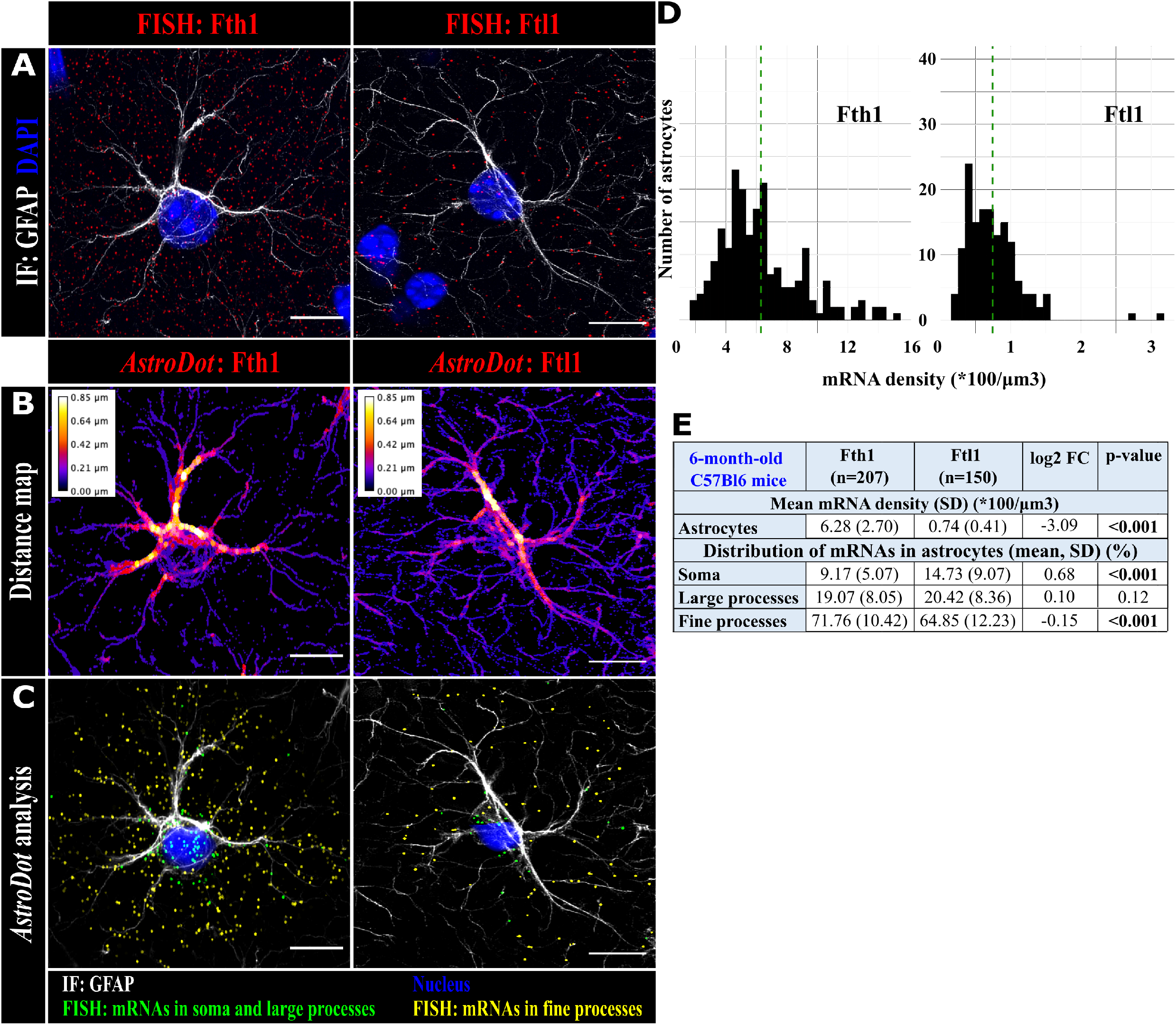
Characterization of ferritin mRNAs level and distribution in 6-month-old hippocampal astrocytes. **(A)** Images of a deconvoluted confocal z-stack of a CA1 astrocyte, with detection by FISH of Fth1 and Ftl1 mRNAs (in red) and detection by co-immunofluorescence (IF) of GFAP (in grey). The nucleus was stained with DAPI (in blue). **(B)** *AstroDot* analysis: Heat map where GFAP-positive process diameter is color-coded. **(C)** *AstroDot* analysis: Each dot corresponds to one FISH dot. Green dots are located in the soma or on GFAP+ large processes (diameter > 0.3 µm); yellow dots are located on GFAP+ fine processes (diameter < 0.3 µm). **(D)** Histogram representation of Fth1 and Ftl1 mRNA density in WT astrocytes (means in dotted lines). **(E)** Table of *AstroDot* calculated mRNA density and distribution. n is the number of cells analyzed. FC means fold change. Statistical significance was determined in Student’s t-test for normally distributed data and equal variances; Welch-Satterthwaite’s test for a normal data distribution and unequal variances; Wilcoxon’s test for non-normally distributed data. Scale bar: 10 µm.

### Ageing increases the Fth1/Ftl1 ratio in hippocampal astrocytic fine processes

Iron accumulates in the brain of healthy subjects with age (Ijomone, Ifenatuoha, Aluko, Ijomone, & Aschner, 2020). Here, we performed FISH experiments on 10-month-old WT mice and compared the level and distribution of ferritin mRNAs between 6- and 10-month-old WT hippocampal astrocytes (**Fig. 2A**). As in 6-month-old mice, Fth1 was expressed at higher levels than Ftl1 (**Fig. 2C, D**). Performing *AstroDo*t analysis, we first observed that the astrocyte volume was identical between both stages (**Fig. S2**). The astrocytic density of Fth1 mRNAs at 10 months was about 1.4-fold significantly higher than at 6 months, while it decreased by 0.8 for Ftl1 in 10-month-old mice (**Fig. 2**). This led to an Fth1/Ftl1 ratio of about 15.2 in 10-month-old mice **(Table 1**). Interestingly, the distribution of Fth1 mRNAs also changed upon ageing **(Fig. 2D)**. We detected higher levels in fine processes and lower levels in large processes. The distribution of Ftl1 mRNAs was unchanged **(Fig. 2D)**. In conclusion, aged WT mice display increased Fth1/Flt1 ratio by about 1.8 in astrocytes **(Table 1)**, and Fth1 mRNAs tend to redistribute in fine astrocytic processes.

**Table 1.** Comparison of ferritin mRNA density and distribution among the 3 conditions analyzed: Ageing, hemochromatosis, Alzheimer’s disease. Means (SD) of *AstroDot* analyses are showed with colors indicating changes (increase in red, decrease in blue). ***, p ≤ 0.001.

| Conditions | mRNA density in astrocytes (*100/$\mu\text{m}^3$) | | | Distribution of mRNA in astrocytes |
| --- | --- | --- | --- | --- |
|  | Fth1 mean (SD) | Ftl1 mean (SD) | Fth1/Ftl1 ratio |  |
| **6-month-old WT** | 6.28 (2.70) | 0.74 (0.41) | 8.49 | Mainly in astrocyte fine processes |
| **Ageing**<br>(6 /10-month-old WT) | 8.95 (3.52)<br>*** | 0.59 (0.21)<br>*** | 15.17 | Fth1 more abundant in fine processes |
| **Hemochromatosis**<br>(6-month-old *Hepc*<sup>-/-</sup> / WT) | 11.24 (5.78)<br>*** | 1.08 (0.41)<br>*** | 10.40 | Unchanged |
| **Alzheimer's disease**<br>(10-month-old APP/PS1dE9 / WT) | 5.88 (2.29)<br>*** | 0.72 (0.29)<br>*** | 8.17 | Fth1 more somatic |

**Figure 2.**
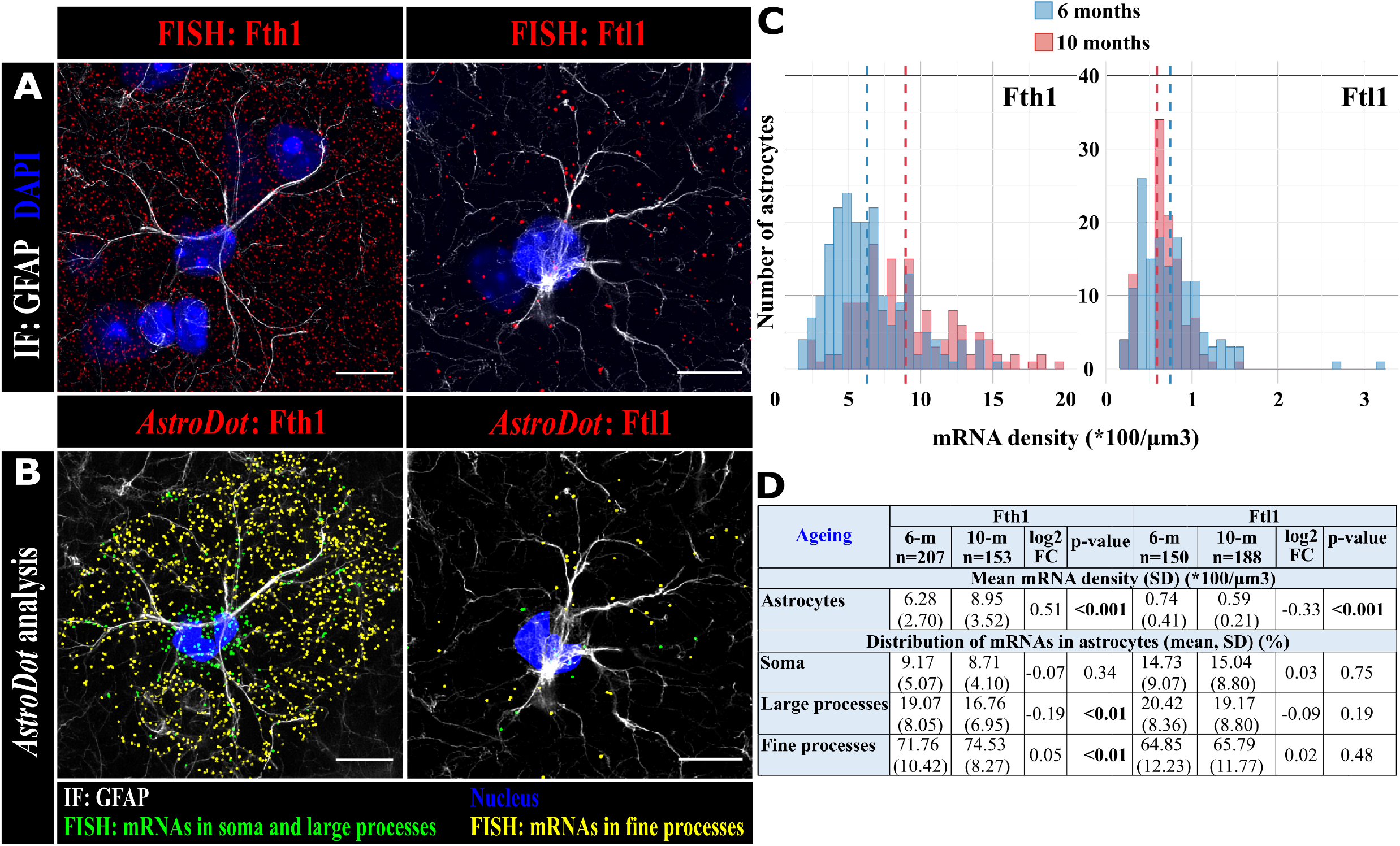
Characterization of ferritin mRNAs level and distribution in ageing hippocampal astrocytes. **A**. Images of a deconvoluted confocal z-stack of a 10-month-old WT CA1 astrocyte, with detection by FISH of Fth1 and Ftl1 mRNAs (in red) and detection by co-immunofluorescence (IF) of GFAP (in grey). The nucleus was stained with DAPI (in blue). **B**. *AstroDot* analysis: Each dot corresponds to one FISH dot. Green dots are located in the soma or GFAP-immunolabelled large processes (diameter > 0.3 µm); yellow dots are located in GFAP-immunolabelled fine processes (diameter < 0.3 µm). **C**. Histogram representation of Fth1 and Ftl1 mRNA density changes in astrocytes upon ageing (means in dotted lines). **D**. Table of *AstroDot* calculated mRNA density and distribution in CA1 astrocytes in 10-month-old mice and comparison with 6-month-old mice. n is the number of cells analyzed. FC means fold change. Statistical significance was determined in Student’s t-test for normally distributed data and equal variances; Welch-Satterthwaite’s test for a normal data distribution and unequal variances; Wilcoxon’s test for non-normally distributed data. Scale bar: 10 µm.

### Hepcidin deletion upregulates ferritin mRNAs and increases the Fth1/Ftl1 ratio in hippocampal astrocytes

Hepcidin, the iron regulatory hormone is mainly expressed by the liver and affects systemic iron availability by controlling ferroportin activity (Atanasiu et al., 2007; Nemeth & Ganz, 2006). Recent data suggest that extra-hepatic sites of hepcidin expression, such as the brain, may be critical in regulating local iron homeostasis (Vujic, 2014). Hepcidin inactivation in the mouse, as well as mutations in Humans, or mutations in proteins regulating hepcidin expression, causes hemochromatosis with a massive and multivisceral iron overload and abnormally high ferritin levels (Brissot et al., 2018; Lesbordes-Brion et al., 2006). Here, we compared the hippocampal astrocytic density and distribution of ferritin mRNAs in 6-month-old WT and hepcidin-deleted mice (*Hepc1*-/-) (Lesbordes-Brion et al., 2006) (**Fig. 3**). Prior to this analysis, we determined that iron level in the brain was higher in *Hepc1*-/- (77.18 ± 21.20) compared to WT (25.69 ± 3.60) (**Fig. 3A**). We observed a smaller volume of astrocytes in *Hepc1*-/- (**Fig. S2**). Interestingly, the astrocytic density of both mRNAs was higher in *Hepc1*-/- mice (**Fig. 3B-E**). The increase of Fth1 (about 1.8-fold) was more pronounced than Ftl1 (about 1.5), which resulted in a higher Fth1/Ftl1 ratio in *Hepc1*-/- mice (**Fig. 3E**) **(Table 1)**. However, mRNA distribution in the soma and fine or large astrocytic processes was unchanged (**Fig. 3E**). These results indicate that Fth1 and Ftl1 mRNAs are both upregulated in hippocampal astrocytes in response to hemochromatosis with no effect on their distribution.

**Figure 3.**
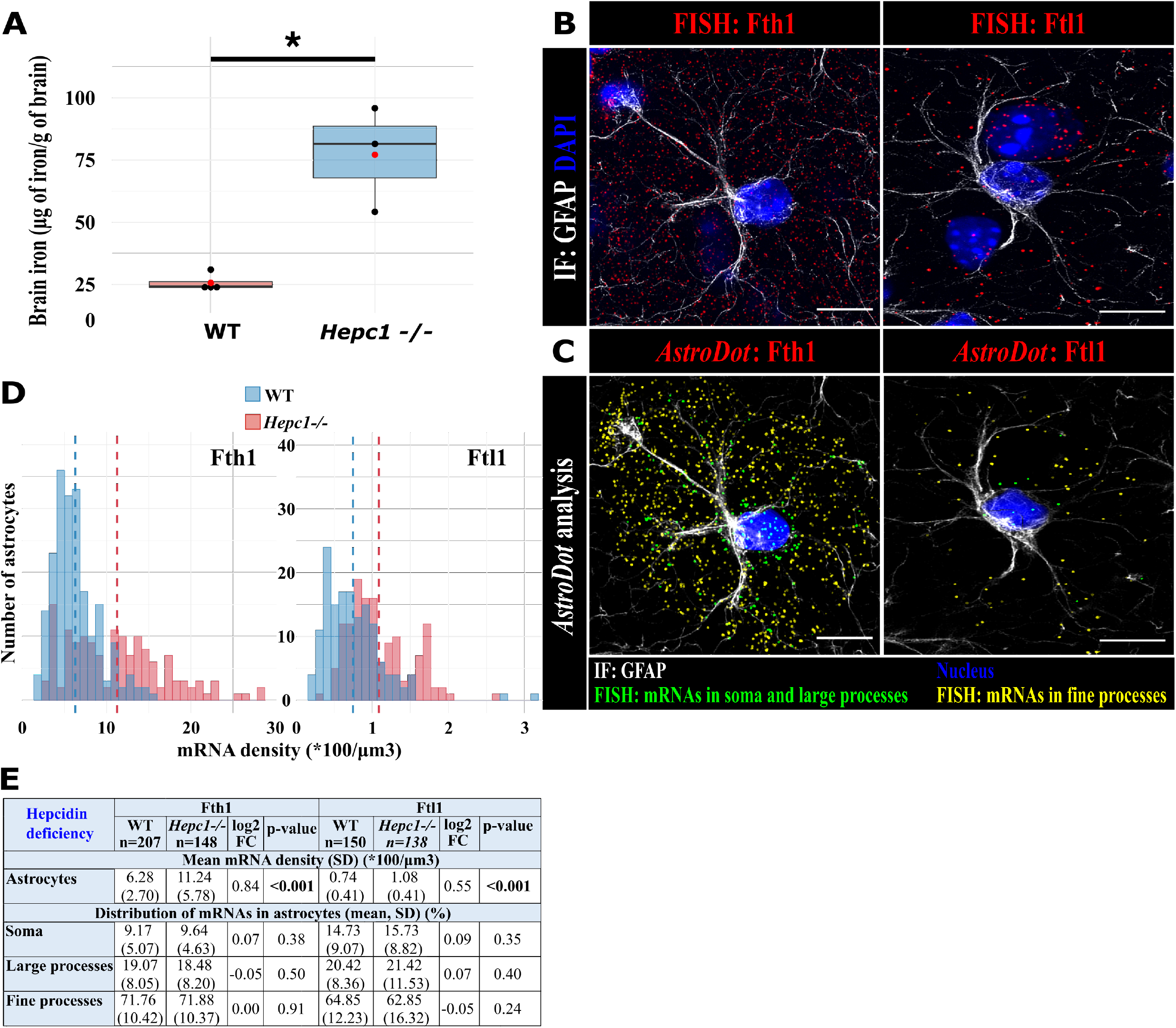
Characterization of ferritin mRNAs level and distribution in hepcidin-deleted CA1 hippocampal astrocytes. **A**. Iron level in the brain of 6-month-old WT and *Hepc-/-* mice (n = 4 WT mice; n= 3 *Hepc-/-; p=0*.*286* Man Whitney one tailed test. **B**. Images of a deconvoluted confocal z-stack of a 6-month-old *Hepc-/-* CA1 astrocyte, with detection by FISH of Fth1 and Ftl1 mRNAs (in red) and detection by co-immunofluorescence (IF) of GFAP (in grey). The nucleus was stained with DAPI (in blue). **C**. *AstroDot* analysis: Each dot corresponds to one FISH dot. Green dots are located in the soma or GFAP-immunolabelled large processes (diameter > 0.3 µm); yellow dots are located in GFAP-immunolabelled fine processes (diameter < 0.3 µm). **D**. Histogram representation of Fth1 and Ftl1 mRNA density changes in astrocytes upon hepcidin deletion (means in dotted lines). Controls are 6-month-old WT mice. **E**. Table of *AstroDot* calculated mRNA density and distribution in CA1 astrocytes in 6-month-old WT and *Hepc-/-* mice and comparison. n is the number of cells analyzed. FC means fold change. Statistical significance was determined in Student’s t-test for normally distributed data and equal variances; Welch-Satterthwaite’s test for a normal data distribution and unequal variances; Wilcoxon’s test for non-normally distributed data. Scale bar: 10 µm.

### Fth1/Ftl1 mRNA ratio is lower and Fth1 mRNAs are redistributed in astrocytic soma in a mouse model of Alzheimer disease

Aberrant iron deposition in the brain has been reported in a number of neurological disorders including AD (Crichton, Dexter, & Ward, 2011; Oshiro, Morioka, & Kikuchi, 2011). Here, we compared the density and localization of ferritin mRNAs in 10-month-old WT and APP/PS1dE9 mice (**Fig. 4**). Amyloid beta (Aß) deposits in the brain parenchyma of APP/PS1dE9 provokes the clustering and strong morphological changes of astrocytes (Escartin et al., 2021). To avoid confounding effects due to Aß-linked astrocytic reactivity, we focused our study on astrocytes not contacting Aß plaques and with a volume comparable to WT (**Fig. S2**). Interestingly, in APP/PS1dE9 mice, we observed a lower Fth1 mRNA astrocytic density (about 1.5-fold), and a higher Ftl1 mRNA density (about 1.2-fold) (**Fig 4C, D**). The distribution of Fth1 mRNAs in APP/PS1dE9 was higher in the soma and lower in astrocytic processes compared to WT mice (**Fig. 4D**). In contrast, the distribution of Ftl1 mRNAs was unchanged (**Fig. 4D**). These results indicate that in APP/PS1dE9 astrocytes, the Fth1/Ftl1 ratio is lower and Fth1 mRNAs become more somatic **(Table 1)**.

**Fig. 4.**
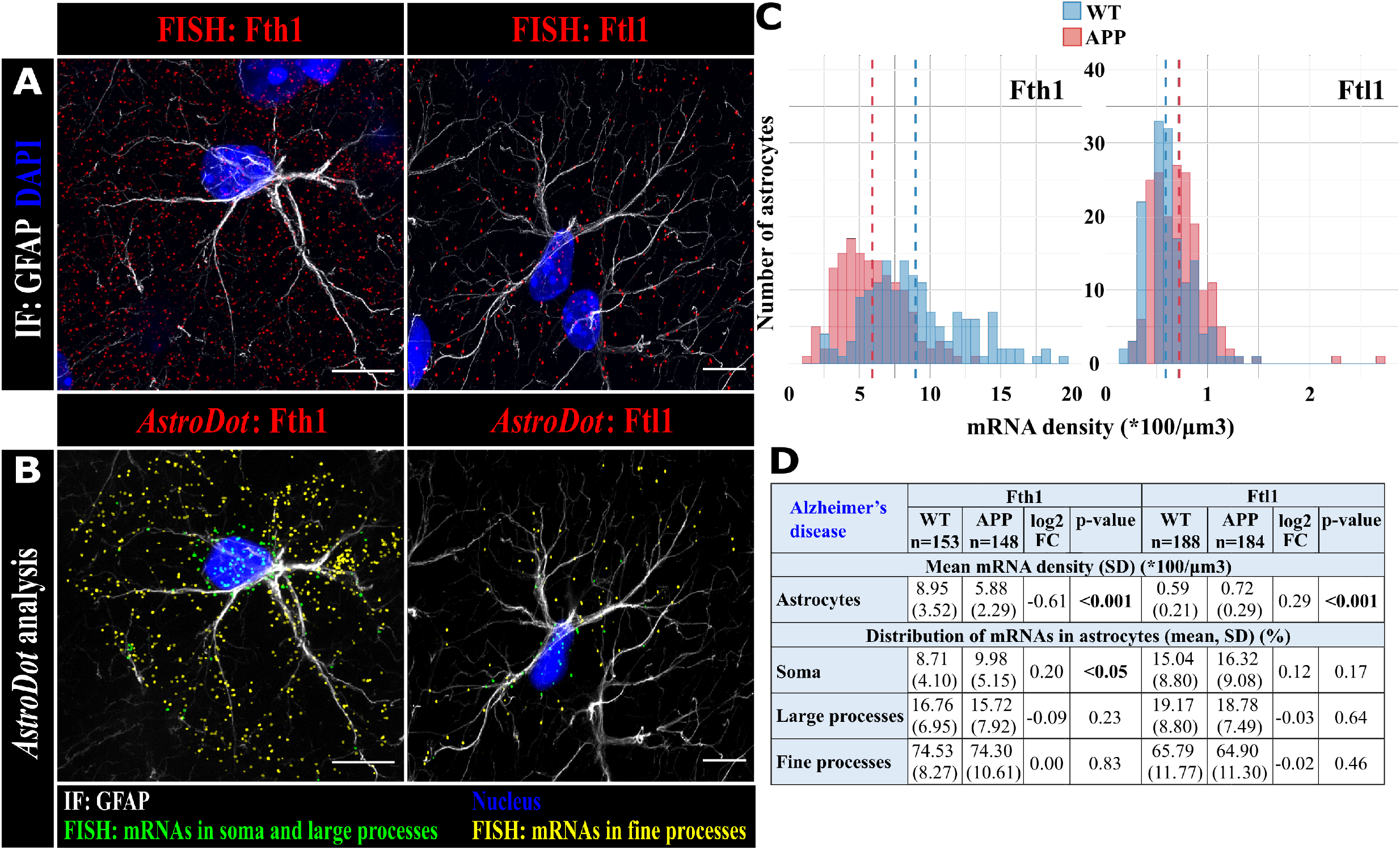
Characterization of Ferritin mRNAs level and distribution in APP/PS1dE9 CA1 hippocampal astrocytes. **A**. Images of a deconvoluted confocal z-stack of a 10-month-old APP/PS1dE9 CA1 astrocyte, with detection by FISH of Fth1 and Ftl1 mRNAs (in red) and detection by co-immunofluorescence (IF) of GFAP (in grey). The nucleus was stained with DAPI (in blue). **B**. *AstroDot* analysis. Each dot corresponds to one FISH dot. Green dots are located in the soma or GFAP-immunolabelled large processes (diameter > 0.3 µm); yellow dots are located in GFAP-immunolabelled fine processes (diameter < 0.3 µm). **C**. Histogram representation of Fth1 and Ftl1 mRNA density changes in astrocytes in APP/PS1dE9 CA1 astrocytes (means in dotted lines). Controls are 10-month-old WT mice. **D**. Table of *AstroDot* calculated mRNA density and distribution in CA1 astrocytes in 10-month-old WT and APP/PS1dE9 mice and comparison. n is the number of cells analyzed. Statistical significance was determined in Student’s t-test for normally distributed data and equal variances; Welch-Satterthwaite’s test for a normal data distribution and unequal variances; Wilcoxon’s test for non-normally distributed data. Scale bar: 10 µm.

## Discussion

Astrocytes are important glial cells which play a pivotal role in iron homeostasis (Dringen et al., 2007). In a previous transcriptomic study, we showed that mRNAs encoding the ferritin light and heavy chains were more present in hippocampal PAPs than in astrocytic soma (Mazare et al., 2020). The same distribution was also found in PvAPs, with however a lower density of Fth1 mRNAs compared to PAPs (Mazare et al., 2020). These data suggested that hippocampal astrocytes express high amount of ferritin and that ferritin would be more translated in astrocytic processes. Here, we used a recently developed *in situ* 3D methodology to analyze ferritin mRNA distribution in astrocytes. One of the main limitations of this method is that only mRNAs located at the level of GFAP+ processes are considered, which, in the case of ferritin, represented about half of mRNAs detected by FISH (**Fig. S1**). Another limitation is the fact that GFAP is less present in extremely fine distal astrocyte processes such as PAPs. Consequently, the localization of non-astrocyte specific mRNAs such as ferritin mRNAs cannot be fully assessed by *AstroDot* (Oudart et al., 2020). Nevertheless, our present results showed that ferritin mRNAs are mostly distributed in astrocytic processes which validates our previous transcriptomic data (Mazare et al., 2020). These data suggest that, in astrocytes, iron buffering and storage mediated by ferritin could be sustained by local translation and would be prevalent in distal compartments.

Ferritin is a 24-mer spherical cage composed by H and L chains. H chains have a ferroxidase center catalyzing the oxidation of ferrous (Fe^2+^) to ferric iron (Fe^3+^), whereas L chains nucleate the mineral core within the ferritin cage. Thus, L-rich ferritins are associated with iron storage, whereas H-rich ferritins are associated with responses to oxidative stress. Interestingly, the composition in ferritin chains varies among tissues (Finazzi & Arosio, 2014). Tissues with a main storage function such as the liver and spleen contain up to 90 % L chains. In contrast, those with high iron oxidation activity such as the heart and brain favor H-rich ferritins (Harrison & Arosio, 1996; Wagstaff, Worwood, & Jacobs, 1978). Our data show a prevalent expression of Fth1 over Ftl1 in astrocytes, indicating that in these cells, ferritins are highly antioxidative. The enrichment of ferritin in astrocyte processes also suggests that ferritin could protect synapse environment from oxidative stress (Codazzi, Pelizzoni, Zacchetti, & Grohovaz, 2015; Dringen et al., 2007).

Production of ferritin upon changes in iron status (overload or depletion) has been shown to be tightly regulated at the transcriptional and post-transcriptional levels by iron regulatory proteins (IRP) binding iron regulatory elements (IRE) on mRNA untranslated regions (Hentze et al., 2010). The regulation by IRPs is exclusively post-transcriptional, it concerns either the translation of mRNAs, this is the case for ferritin which has its IRE in 5’, or the stability of mRNA when the IRE is in 3’. The transcriptional level of ferritin regulation is predominantly achieved by the transcription factor NRF2 which is a key antioxidant actor in the case of iron overload (Song, Gao, Sheng, Rui, & Luo, 2021). Here, our analysis in aging, hemochromatosis, and Alzheimer’s disease mouse models showed first that mRNA density varies in astrocytes in all conditions (**Table 1**). Fth1 was upregulated in ageing and hepcidin-deleted mice and downregulated in APP/PS1dE9 mice, while Ftl1 was upregulated in hepcidin-deleted and APP/PS1dE9 mice and downregulated in ageing mice. Second, our results showed variation of the Fth1/Flt1 ratio among the tested conditions. It was higher in ageing and hepcidin-deleted mice compared to controls. These results suggest that the antioxidant effect of ferritin might increase and protect the brain. In great contrast to ageing and hepcidin-deleted mice, we found a lower Fth1/Ftl1 ratio in APP/PS1dE9 astrocytes. AD has been linked to oxidative stress with imbalanced free radical production and antioxidant defenses (Belaidi & Bush, 2016; Liu et al., 2019; Molina-Holgado et al., 2007; Zecca et al., 2004). The upregulation of Ftl1 and the downregulation of Fth1 leading to a lower Fth1/Ftl1 ratio in APP/PS1dE9 astrocytes could likely be deleterious by increasing the risk of oxidative stress and exacerbating brain tissue damages. Finally, Fth1 mRNAs became more somatic in APP/PS1dE9 mice. This result contrast with ageing and hepcidin-deleted mice where ferritin mRNA preferential distribution in fine astrocytic processes was mostly unchanged. It suggests that in AD iron dysregulation may originate from altered distribution of ferritin mRNAs in astrocytes.

To conclude, our study indicates that ferritin transcription, mRNA distribution, and regulation of Fth1/Ftl1 ratio in astrocytes is context-dependent and might contribute to iron dys-homeostasis in physiopathology.

## Acknowledgements

This work was funded by the Fondation pour la Recherche Médicale to M. Cohen-Salmon (AJE20171039094) and N. Mazaré (FDT201904008077), the Fondation France Alzheimer to M. Cohen-Salmon and Carole Escartin, the ED3C doctoral school to N. Mazaré and M. Oudart and the “Journées de Neurologie de Langue Française” to R. Tortuyaux. The creation of the Center for Interdisciplinary Research in Biology (CIRB) was funded by the “Fondation Bettencourt Schueller”. C. Escartin’s work was funded by the French national research agency ANR-16-TERC-0016-01.

## References

Abbott, N. J., Ronnback, L., & Hansson, E. (2006). Astrocyte-endothelial interactions at the blood-brain barrier. Nat Rev Neurosci, 7 (1), 41–53. Retrieved from http://www.ncbi.nlm.nih.gov/entrez/query.fcgi?cmd=Retrieve&db=PubMed&dopt=Citation&list_uids=16371949

Alvarez, J. I., Katayama, T., & Prat, A. (2013). Glial influence on the blood brain barrier. Glia. doi:10.1002/glia.22575

Atanasiu, V., Manolescu, B., & Stoian, I. (2007). Hepcidin--central regulator of iron metabolism. Eur J Haematol, 78 (1), 1–10. doi:10.1111/j.1600-0609.2006.00772.x

Belaidi, A. A., & Bush, A. I. (2016). Iron neurochemistry in Alzheimer’s disease and Parkinson’s disease: targets for therapeutics. J Neurochem, 139 Suppl 1, 179–197. doi:10.1111/jnc.13425

Brissot, P., Pietrangelo, A., Adams, P. C., de Graaff, B., McLaren, C. E., & Loreal, O. (2018). Haemochromatosis. Nat Rev Dis Primers, 4, 18016. doi:10.1038/nrdp.2018.16

Cheli, V. T., Correale, J., Paez, P. M., & Pasquini, J. M. (2020). Iron Metabolism in Oligodendrocytes and Astrocytes, Implications for Myelination and Remyelination. ASN Neuro, 12, 1759091420962681. doi:10.1177/1759091420962681

Cheli, V. T., Santiago Gonzalez, D. A., Wan, Q., Denaroso, G., Wan, R., Rosenblum, S. L., & Paez, P. M. (2021). H-ferritin expression in astrocytes is necessary for proper oligodendrocyte development and myelination. Glia, 69 (12), 2981–2998. doi:10.1002/glia.24083

Codazzi, F., Pelizzoni, I., Zacchetti, D., & Grohovaz, F. (2015). Iron entry in neurons and astrocytes: a link with synaptic activity. Front Mol Neurosci, 8, 18. doi:10.3389/fnmol.2015.00018

Cohen-Salmon, M., Slaoui, L., Mazare, N., Gilbert, A., Oudart, M., Alvear-Perez, R., … Boulay, A. C. (2021). Astrocytes in the regulation of cerebrovascular functions. Glia, 69 (4), 817–841. doi:10.1002/glia.23924

Connor, J. R., & Menzies, S. L. (1995). Cellular management of iron in the brain. J Neurol Sci, 134 Suppl, 33–44. doi:10.1016/0022-510x(95)00206-h

Crichton, R. R., Dexter, D. T., & Ward, R. J. (2011). Brain iron metabolism and its perturbation in neurological diseases. J Neural Transm (Vienna), 118 (3), 301–314. doi:10.1007/s00702-010-0470-z

Dallerac, G., Zapata, J., & Rouach, N. (2018). Versatile control of synaptic circuits by astrocytes: where, when and how? Nat Rev Neurosci, 19 (12), 729–743. doi:10.1038/s41583-018-0080-6

Dringen, R., Bishop, G. M., Koeppe, M., Dang, T. N., & Robinson, S. R. (2007). The pivotal role of astrocytes in the metabolism of iron in the brain. Neurochem Res, 32 (11), 1884–1890. doi:10.1007/s11064-007-9375-0

Escartin, C., Galea, E., Lakatos, A., O’Callaghan, J. P., Petzold, G. C., Serrano-Pozo, A., … Verkhratsky, A. (2021). Reactive astrocyte nomenclature, definitions, and future directions. Nat Neurosci, 24 (3), 312–325. doi:10.1038/s41593-020-00783-4

Finazzi, D., & Arosio, P. (2014). Biology of ferritin in mammals: an update on iron storage, oxidative damage and neurodegeneration. Arch Toxicol, 88 (10), 1787–1802. doi:10.1007/s00204-014-1329-0

Harrison, P. M., & Arosio, P. (1996). The ferritins: molecular properties, iron storage function and cellular regulation. Biochim Biophys Acta, 1275 (3), 161–203. doi:10.1016/0005-2728(96)00022-9

Hentze, M. W., Muckenthaler, M. U., Galy, B., & Camaschella, C. (2010). Two to tango: regulation of Mammalian iron metabolism. Cell, 142 (1), 24–38. doi:10.1016/j.cell.2010.06.028

Hohnholt, M. C., & Dringen, R. (2013). Uptake and metabolism of iron and iron oxide nanoparticles in brain astrocytes. Biochem Soc Trans, 41 (6), 1588–1592. doi:10.1042/BST20130114

Ijomone, O. M., Ifenatuoha, C. W., Aluko, O. M., Ijomone, O. K., & Aschner, M. (2020). The aging brain: impact of heavy metal neurotoxicity. Crit Rev Toxicol, 50 (9), 801–814. doi:10.1080/10408444.2020.1838441

Jankowsky, J. L., Fadale, D. J., Anderson, J., Xu, G. M., Gonzales, V., Jenkins, N. A., … Borchelt, D. R. (2004). Mutant presenilins specifically elevate the levels of the 42 residue beta-amyloid peptide in vivo: evidence for augmentation of a 42-specific gamma secretase. Hum Mol Genet, 13 (2), 159–170. doi:10.1093/hmg/ddh019

Lesbordes-Brion, J. C., Viatte, L., Bennoun, M., Lou, D. Q., Ramey, G., Houbron, C., … Vaulont, S. (2006). Targeted disruption of the hepcidin 1 gene results in severe hemochromatosis. Blood, 108 (4), 1402–1405. doi:10.1182/blood-2006-02-003376

Liu, C., Liang, M. C., & Soong, T. W. (2019). Nitric Oxide, Iron and Neurodegeneration. Front Neurosci, 13, 114. doi:10.3389/fnins.2019.00114

Lozoff, B., Beard, J., Connor, J., Barbara, F., Georgieff, M., & Schallert, T. (2006). Long-lasting neural and behavioral effects of iron deficiency in infancy. Nutr Rev, 64(5 Pt 2), pS34-43; discussion S72-91. doi:10.1301/nr.2006.may.s34-s43

Mazare, N., Oudart, M., Moulard, J., Cheung, G., Tortuyaux, R., Mailly, P., … Cohen-Salmon, M. (2020). Local Translation in Perisynaptic Astrocytic Processes Is Specific and Changes after Fear Conditioning. Cell Reports, 32 (8), 108076. doi:10.1016/j.celrep.2020.108076

Molina-Holgado, F., Hider, R. C., Gaeta, A., Williams, R., & Francis, P. (2007). Metals ions and neurodegeneration. Biometals, 20(3-4), 639–654. doi:10.1007/s10534-006-9033-z

Nemeth, E., & Ganz, T. (2006). Regulation of iron metabolism by hepcidin. Annu Rev Nutr, 26, 323–342. doi:10.1146/annurev.nutr.26.061505.111303

Oshiro, S., Morioka, M. S., & Kikuchi, M. (2011). Dysregulation of iron metabolism in Alzheimer’s disease, Parkinson’s disease, and amyotrophic lateral sclerosis. Adv Pharmacol Sci, 2011, 378278. doi:10.1155/2011/378278

Oudart, M., Tortuyaux, R., Mailly, P., Mazare, N., Boulay, A. C., & Cohen-Salmon, M. (2020). AstroDot - a new method for studying the spatial distribution of mRNA in astrocytes. J Cell Sci, 133 (7). doi:10.1242/jcs.239756

Perea, G., Navarrete, M., & Araque, A. (2009). Tripartite synapses: astrocytes process and control synaptic information. Trends Neurosci, 32 (8), 421–431. doi:S0166-2236(09)00101-5 [pii] 10.1016/j.tins.2009.05.001

Raha-Chowdhury, R., Raha, A. A., Forostyak, S., Zhao, J. W., Stott, S. R., & Bomford, A. (2015). Expression and cellular localization of hepcidin mRNA and protein in normal rat brain. BMC Neurosci, 16, 24. doi:10.1186/s12868-015-0161-7

Song, S., Gao, Y., Sheng, Y., Rui, T., & Luo, C. (2021). Targeting NRF2 to suppress ferroptosis in brain injury. Histol Histopathol, 36 (4), 383–397. doi:10.14670/HH-18-286

Stephenson, E., Nathoo, N., Mahjoub, Y., Dunn, J. F., & Yong, V. W. (2014). Iron in multiple sclerosis: roles in neurodegeneration and repair. Nat Rev Neurol, 10 (8), 459–468. doi:10.1038/nrneurol.2014.118

Torrance, J. D., & Bothwell, T. H. (1980). Tissue iron stores (J. D. Cook Ed. Vol. 1). New York.

Vujic, M. (2014). Molecular basis of HFE-hemochromatosis. Front Pharmacol, 5, 42. doi:10.3389/fphar.2014.00042

Wagstaff, M., Worwood, M., & Jacobs, A. (1978). Properties of human tissue isoferritins. Biochem J, 173 (3), 969–977. doi:10.1042/bj1730969

Wang, X. S., Ong, W. Y., & Connor, J. R. (2002). A light and electron microscopic study of divalent metal transporter-1 distribution in the rat hippocampus, after kainate-induced neuronal injury. Exp Neurol, 177 (1), 193–201. doi:10.1006/exnr.2002.7962

Zecca, L., Youdim, M. B., Riederer, P., Connor, J. R., & Crichton, R. R. (2004). Iron, brain ageing and neurodegenerative disorders. Nat Rev Neurosci, 5 (11), 863–873. doi:10.1038/nrn1537

Zhang, X., Gou, Y. J., Zhang, Y., Li, J., Han, K., Xu, Y., … Gao, G. (2020). Hepcidin overexpression in astrocytes alters brain iron metabolism and protects against amyloid-beta induced brain damage in mice. Cell Death Discov, 6 (1), 113. doi:10.1038/s41420-020-00346-3

