## Supplemental Figure S1, S2 and Table S1 for "Physiopathological changes of ferritin mRNA density and distribution in hippocampal astrocytes in the mouse brain"

**Supporting Information (Figure S1, S2 and Table S1)**

**A**

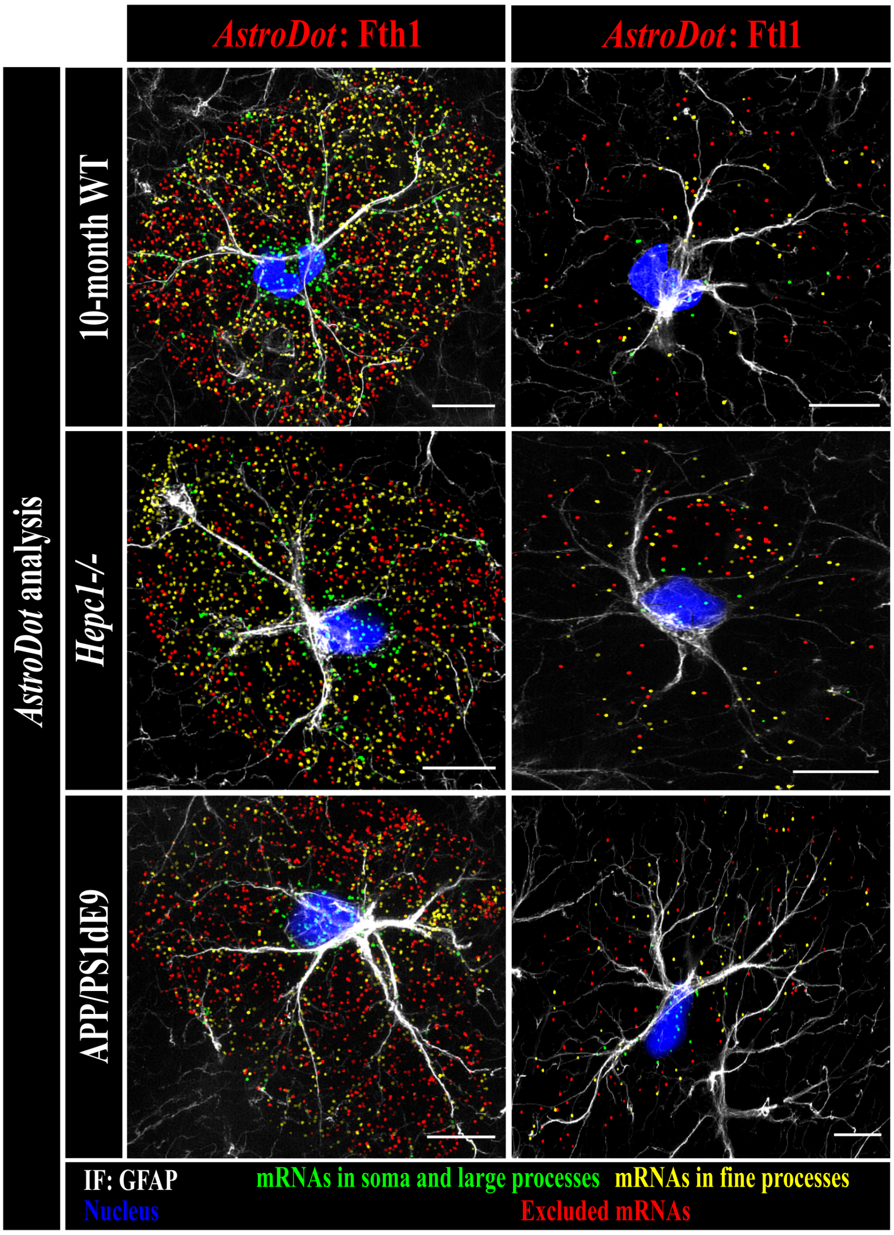


**B**

| Conditions | Fth1 | | Ftl1 | |
|---|---|---|---|---|
| | mean (SD) | p value | mean (SD) | p value |
| **Ageing** (6/10-month-old WT) | 51.98 (13.81) | 0.43 | 45.86 (11.99) | **<0.001** |
| | 50.94 (11.00) | | 53.15 (9.67) | |
| **Hemochromatosis** (6-month-old *Hepc-/-* / WT) | 45.15 (10.68) | **<0.001** | 40.36 (14.98) | **<0.001** |
| **Alzheimer's disease** (10-month-old APP/PS1dE9 / WT) | 52.04 (16.33) | 0.50 | 52.22 (10.73) | 0.38 |

**Fig. S1 AstroDot analysis of ferritin FISH dots not localized on GFAP-immunostaining. A.** *AstroDot* analysis. Each dot corresponds to one FISH dot. Green dots are located in the soma or GFAP-immunolabelled large processes (diameter > 0.3 µm); yellow dots are located in GFAP-immunolabelled fine processes (diameter < 0.3 µm); red dots do not localize on GFAP immunostaining (i.e. excluded mRNAs). **B.** Percentage of ferritin mRNA FISH dots not localized on GFAP-immunostaining according to conditions. Statistical significance was determined in Student's t-test for normally distributed data and equal variances; Welch-Satterthwaite's test for a normal data distribution and unequal variances; Wilcoxon's test for non-normally distributed data.

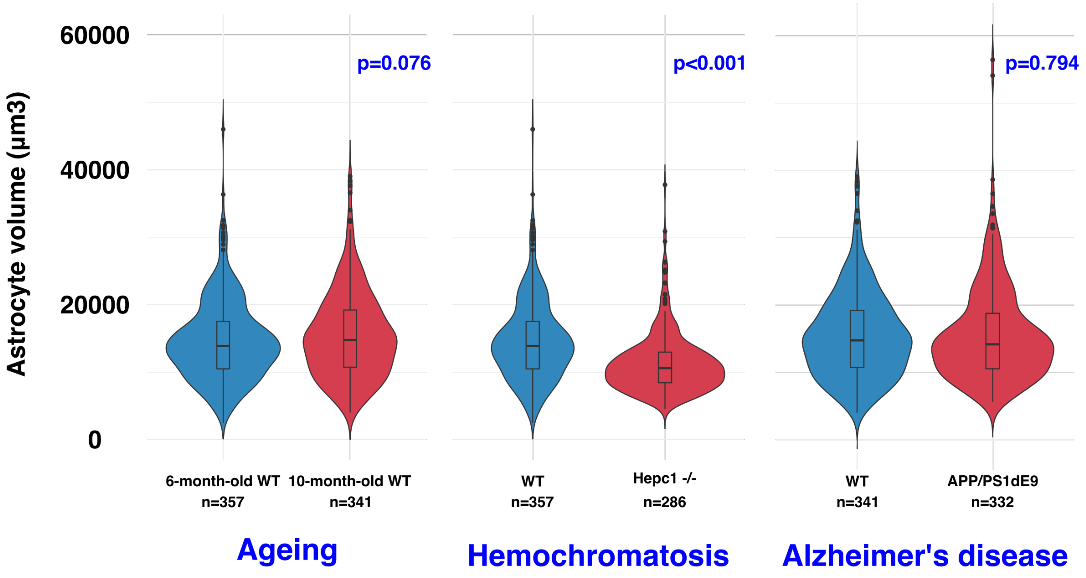


**Fig S2 *AstroDot* estimation of astrocyte volume. A.** Violin plots including a boxplot of astrocyte volumes. n is the number of cells analyzed per condition. Statistical significance was determined by a Student's t-test for normally distributed data and equal variances; Welch-Satterthwaite's test for a normal data distribution and unequal variances; Wilcoxon's test for non-normally distributed data.

| FISH reagents | | |
|---|---|---|
| **Name** | **Supplier** | **Reference** |
| RNAscope® Multiplex Fluorescent detection reagent V2 | Advanced Cell Diagnostic | 323110 |
| RNAscope® $H_2O_2$ and Protease | Advanced Cell Diagnostic | 322381 |
| Fluoromount-G® | Southern Biotech | 0100-01 |
| Coverglass 0.13 – 0.17mm thick | Immuno Cell | 65.300.13 |
| SuperFrost® Plus slides | VWR | 631-0108 |
| Hydrophobic immunostaining pen | Vector laboratories | H-4000 |

| FISH RNA probes | | |
|---|---|---|
| **Gene name** | **Probe** | **Reference and supplier** |
| Fth1 | RNAscope® Probe – Mm-Fth1 | 511561 Advanced Cell Diagnostic |
| Ftl1 | RNAscope® Probe – Mm-Ftl1 | 314771 Advanced Cell Diagnostic |

| FISH Fluorophore | | |
|---|---|---|
| **Name** | **Reference and supplier** | **Dilution** |
| Opal 570 | FP14488A Perkin Elmer | 1:1500 |
| Opal 650 | FP1496A Perkin Elmer | 1:1500 |

| Antibodies | | |
|---|---|---|
| **Name** | **Reference and supplier** | **Dilution** |
| Rabbit anti-Glial Fibrillary Acidic Protein (GFAP) antibody | G9269 Sigma | 1:500 |
| Goat anti-Rabbit IgG (H+L) Highly Cross-Adsorbed Secondary Antibody, Alexa Fluor 488 | A11034 Invitrogen | 1:1500 |

**Table S1: Key resources**
